# Regulating lipid composition rationalizes acyl-tail saturation homeostasis in ectotherms

**DOI:** 10.1101/870063

**Authors:** M. Girard, T. Bereau

## Abstract

Cell membranes mainly consist of lipid bilayers with an actively regulated composition. The underlying processes are still poorly understood, in particular how the hundreds of components are controlled. Surprisingly, in recent experiments on ectotherms, the cholesterol fraction, along with un- and mono-saturated acyl tail fractions and demixing temperatures, was shown to increase with body temperature. We establish a model based on chemical reaction networks to study regulation of membranes, resulting in multiple semi-grand canonical ensembles. By running computer simulations, we show that higher cholesterol fractions correlate with lower degrees of unsaturation, ultimately controlling the composition of lipid tails. Cholesterol also dictates membrane viscosity and regulation of the latter implies that cholesterol must increase with temperature. Overall, our model proposes a different picture of lipid regulation, where components can be passively, instead of actively, regulated.

**SIGNIFICANCE:** In this article, we propose a regulation model where only some of the components are actively regulated between membranes, while others are naturally balanced by chemical potentials. This model provides a rationale to recently measured puzzling trends in ectotherms, that is, increased plasma membrane cholesterol fraction with temperature. Here, we show that it is directly correlated with with acyl tail saturation and order parameter correlation length. Furthermore, we highlight the relation between cholesterol and membrane viscosity.

## BACKGROUND

For eukaryotes, the plasma membrane is the last interface between the cell interior and the extracellular environment. It is responsible for regulating permeation of molecules (1, 2), and membrane protein function (3, 4). Cell membranes mainly consist of lipid bilayers. In aqueous environments, these amphiphilic molecules, comprised of a polar headgroup and hydrocarbon tails, readily self-assemble to hydrate the polar head while minimizing hydrophobic interactions of apolar tails. Lipids can be broadly classified by their headgroup (phosphatidylcholine, phosphatidylserine, etc.). Most lipids in biological membranes have two hydrocarbon tails, and their nature is generally widely variable, ranging from 12 to 24 carbons with varying degrees of saturation. When considering both classifications, this yields hundreds to thousands of different lipid types (5). Furthermore, biological membranes are generally asymmetric: the two leaflets of the bilayers have different compositions, which is maintained by transmembrane proteins pumping specific lipids from one leaflet to the other (6). Similar complexity is replicated by various lipid membranes inside the cell and, while advances have been made by lipidomics, many fundamental questions still need to be answered (7).

Membrane regulation is one of the most poorly understood areas of biological membrane physics. The question of how and why various properties are regulated is of great interest to biology. For instance, there are differences in composition of various membranes within single cells that are thought to endow the membrane with specific properties (8). Lipid tail composition is often thought to be regulated by specific pathways activated by sensor proteins in the membrane, but little is known about them (9). Based on experimental observations, two physical properties are perceived as tightly regulated in plasma membranes: viscosity (10, 11) and curvature (12), which correlate *in-vivo* (13). Regulation of viscosity includes acyl tail remodeling via the Lands cycle (14–16), a process through which lipids change the nature of their acyl tails. Membrane curvature regulation involves multiple factors. Some of them are internal, for instance phosphatidylethanolamine tends to produce negative curvature (17). However, it may also involve external factors to the membrane, such as action of the cytoskeleton (18). Another quantity has recently emerged as being regulated in cells: the difference between ambient and bilayer demixing temperature (*T*_m_). Whether *T*_m_ is regulated through the same mechanism as viscosity or simply as a byproduct of its regulation is currently unknown. Additionally, while there is a clear link between *T*_m_ and proximity to the critical temperature needed for lipid rafts, the picture remains incomplete. However, experiments presented in (19) clearly demonstrate that lowering zebrafish body temperature by ≈ 12 K lowers the *T*_m_ of their plasma membranes by ≈ 12 K, an astounding precision considering how much the membrane composition changes in the process. Surprisingly, the cholesterol fraction in zebrafish *increases* with temperature. In model membranes, increasing cholesterol is linked with lower transition temperatures (20).

In order to simulate realistic biological behavior on the computer, the composition must be allowed to fluctuate. This is elegantly achieved by controlling the chemical potential—the thermodynamic variable conjugate to composition. Alternatively, differences between chemical potentials of different chemical species can be fixed, yielding the semi-grand canonical (SGC) ensemble. This keeps the overall number of molecules fixed, a more suitable approach for molecular dynamics, as compared to the grand canonical ensemble. Furthermore, chemical reaction networks, sets of reactants, products and reactions, can be expressed into chemical potential differences (21) between molecules and large-scale, parallel algorithms have been developed for this ensemble (22).

Here, we model the regulation pathway of phospholipids as a chemical network, creating an SGC ensemble per regulated phospholipid type, the collection of which we call the regulated thermodynamic ensemble (RTE). We apply this methodology to a phosphatidylcholine (PC) membrane incorporating various amounts of cholesterol. We show that increasing cholesterol fraction leads to an overall increase in acyl tail saturation and viscosity. This provides a rationale for lipid changes observed in (19): under the homeoviscous hypothesis, cholesterol concentration must increase with temperature in order to compensate for the decrease in viscosity induced by temperature. Furthermore, acyl tail saturation changes are surprisingly similar to those of (19). This also provides an explanation for the correlation between cholesterol enrichment (9) and acyl tail desaturation (10).

## REGULATED ENSEMBLES AS A COLLECTION OF SEMI-GRAND CANONICAL ENSEMBLES

In this section, we introduce a hypothetical regulation pathway to illustrate the collection of SGC ensembles. We make a connection with the biological pathway responsible for remodeling the acyl tail nature, although similar arguments can be applied to other mechanisms involving chemical networks (9). The RTE can be used to describe out-of-equilibrium systems, such as asymmetric membranes, and derived results should be associated with homeostasis or stationary states rather than a thermodynamic equilibrium in the usual sense, since chemical potential differences can be enforced.

Let us consider a simple lipid bilayer, with only two classes of lipids (head groups) and acyl tails of varying saturation, as illustrated in Fig. 1 (A). Inside the cell, lipid transport proteins (LTP) provide most of the trafficking and we neglect vesicular transport (23) for simplicity. LTPs transport lipids from one membrane to another, and we hypothesize that regulatory transport mainly takes place with the endoplasmic reticulum (ER). For simplicity, we also discard effects of other proteins present in biological membranes (24).

**Figure 1:**
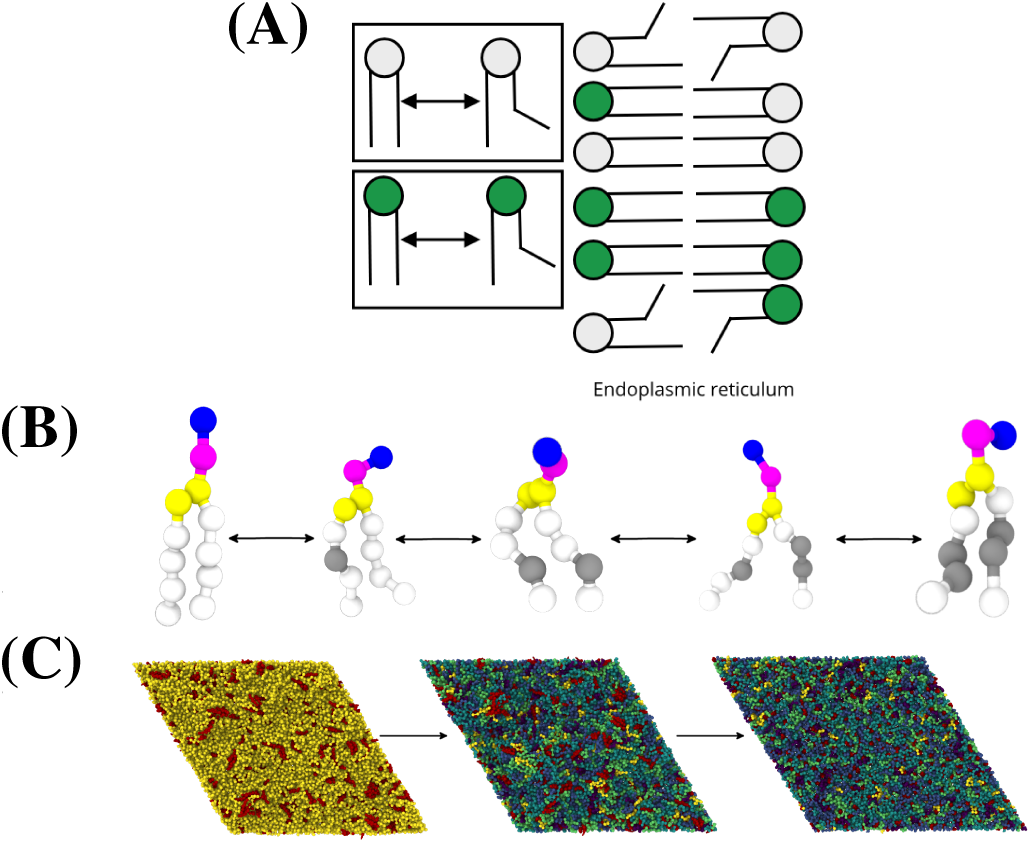
(A) Sketch of the regulatory processes used for our argument. Lipids are moved from the ER to other membranes through LTP. Lipids being able to transform from one to another through the Lands cycle, indicated by arrows, belong to a single SGC, depicted here by rectangles. The remodeling cycle enforces chemical potential differences within the SGC ensemble, which is then carried to other membranes by LTP. Headgroup composition is fixed as head groups are not able to remodel from one to another. (B) Representative ensemble of PC molecules used in molecular dynamics, with different degrees of unsaturation. During simulation, at each Monte-Carlo event, molecules may gain or lose a unsaturated bond. Unsaturations are located on gray beads and only the two middle beads of each chain can be unsaturated. Since there are multiple locations available for unsaturation, many different molecules may have the same degree of unsaturation. (C) Time evolution of a lipid bilayer initialized with two unsaturations per acyl tail. Left: after initial relaxation and thermalization (1ns), middle: after 100 ns, right: after 15 *µ*s. Composition relaxation is faster than cholesterol diffusion and dispersion throughout the layer. Lipids are colored by unsaturations, ranging from zero (blue) to four (yellow), while cholesterol is colored red.

In this picture, lipids present on the plasma membranes are simply a subset of a wider chemical-reaction network, which includes remodeling in ER and transport by LTP. The former implies formation of a protein-lipid complex, which involves at least partial extraction of the lipid, creating a free-energy barrier to form the complex. For a lipid of species *s* in a membrane with a given composition, rates of formation of complexes depend on the free-energy difference of the lipid between barrier and membrane states. Once a lipid molecule is remodeled, membrane composition changes, free energy decreases and remodeling becomes slower. If the system is at equilibrium and closed, then it converges towards the global free-energy minimum (Δ*µ* = 0). Enzymatic pathways are usually described by Michaelis-Menten kinetics (25) where formation of protein-substrate complexes is slow and reversible, while irreversible transformation into products is fast. Biological chemical networks exhibit complex kinetics and are often out-of-equilibrium due to externally controlled co-factors, for instance adenosine triphosphate (ATP) concentration (26–28). Such chemical networks can be described by a non-equilibrium Gibbs free energy (21), which displaces the steady state away from equilibrium.

The Lands cycle, the acyl tail remodeling process, is an interesting chemical network because it involves many lipid species, but few enzymes. These are able to cleave an acyl tail from a PC molecule, yielding a fatty acid and a lysophosphatidylcholine (LPC) or to attach a coenzyme A-fatty acid complex onto an LPC molecule to produce a PC (14, 15, 29, 30). Within this cycle, we can consider transformation from PC to LPC as reactions of the chemical network and co-factors as chemo-statted i.e., externally controlled. This definition is somewhat arbitrary as these components may be regulated themselves. Overall, this enforces a chemical potential difference between all PC and LPC species (Δ*µ* ≠ 0), which is dependent on co-factor concentrations and affinities (i.e., binding rates to different substrates). If multiple chemical networks are isolated as a function of external co-factors, then each network constitutes a single SGC ensemble. Molecules within large SGC ensembles are likely to form complex mixtures with properties responding to other single chemo-statted molecules. As we will show later, experimental observations on zebrafish can be explained if lipid tails respond to cholesterol concentrations.

Enzyme affinities are currently not well characterized. In order to perform simulations however, all values of Δ*µ* between different lipids species must be specified. We therefore make a number of approximations. Since phospholipids such as PC are the main component of membranes, we will assume that the acylation of LPC occurs much faster than the hydrolyzation of PC, or equivalently, that the concentration of the phospholipase is sufficiently low to fully depopulate LPC. Indeed, LPC, and lypophospholipids in general, are minor components of cell membranes. When a PC molecule goes through the cycle—is hydrolyzed and acylated—one of the acyl tails will be replaced by a new one. Here, we assume that phospholipases do not selectively hydrolyze lipids, such that the activated state does not depend on the acyl tail nature, and further that the new acyl tail has a random saturation sampled from a distribution that is at equal chemical potentials. Presumably, this should also entail the regulation of fatty acids, which is outside the scope of this article. Nevertheless, if such conditions are satisfied, then all PC concentrations are at equilibrium such that Δ*µ*_s,s′_ = 0 for any two species *s, s*′, part of the same SGC. We call this the equal-binding approximation. Here, we are interested in lipids with a single acyl tail-length, which are typically transported by a single LTP—devoid of co-factors and thus playing no role in lipid concentrations.

## PHOSPHATIDYLCHOLINES AND CHOLESTEROL IN REGULATED ENSEMBLES

Making use of the equal-binding approximation, we consider a simple RTE lipid membrane consisting of a single SGC ensemble, constituted by 16:(0-2), 16:(0-2) PC molecules (with 0 to 2 unsaturations per acyl tail, see Fig 1 (B, C)) and mixed with cholesterol. Overall, the PC SGC ensemble contains 16 distinct lipid molecules. We consider a mixture of 1600 PC molecules along with a molar fraction of 10 − 30% cholesterol. Systems are kept at constant temperatures between 289 and 314 K (*k*_B_*T* = 2.4 − 2.6 kJ/mol), which results in a homogeneous liquid phase, as shown in Fig 2 (A). The nematic order parameter, characterized by the largest eigenvalue of the de Gennes **Q**-tensor, *λ*_+_, is characteristic of liquid-disordered phases and increases with cholesterol fraction, a known effect in fluid membranes (31).

**Figure 2:**
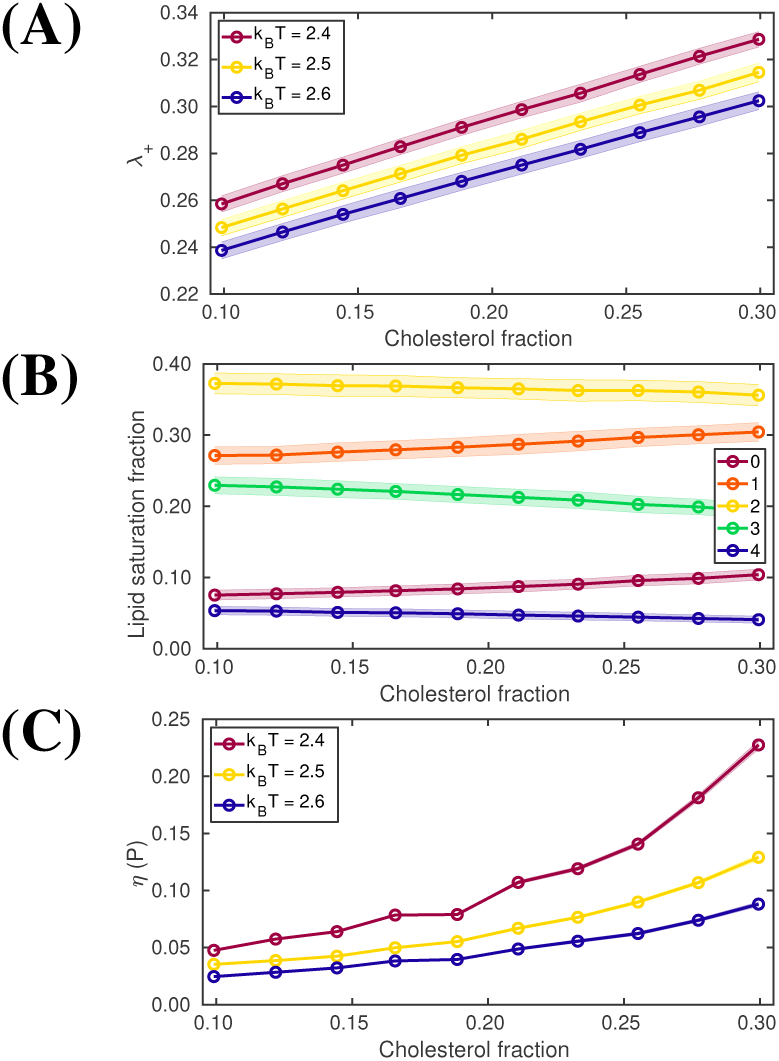
(A) Nematic order of lipid molecules in the system. (B) Lipid tail composition of 16:(0-2), 16:(0-2) PC lipids under the equal binding approximation at *k*_B_*T* = 2.5. Color indicates the overall number of unsaturations on the lipid molecule. (C) Viscosity of the membrane as a function of cholesterol fraction. Shaded area indicates the standard deviation of the data. For viscosity, it is extracted from the confidence interval of the fitting procedure and is of similar width as the curve

Lipid unsaturation fraction, the fraction of lipids in the membrane with a given number of unsaturations, is shown in Fig. 2 (B). It is essentially dominated by the binomial distribution of molecule types associated with saturation degree. For instance, there is only a single fully saturated molecule and a single molecule with four unsaturations, while there are six molecules with two unsaturations. Furthermore, there is a slight bias towards saturated tails. Cholesterol appears as a saturating agent in these membranes, i.e., the proportion of acyl chains with low unsaturation increases with cholesterol. This is associated with a decrease in area per lipid in the membrane (see Fig. S2), meaning that an increase in cholesterol increases packing. Since saturated lipids pack better, their proportion increases and this explains the bias towards saturated tails as cholesterol is added. As a general rule, unsaturations positioned far from the headgroup are preferred by cholesterol. A weak preference for unsaturations to be located on glycol carbon position *sn*1 over *sn*2 is also observed. Characterizing the correlation of the nematic order parameter in the RTE shows short (sub-nm, see Fig S1) correlation lengths, which increase with cholesterol and decrease with temperature, similar to *λ*_+_, and that persist when the membrane composition is fixed at a lower temperature. This would suggest that *T*_m_ increases with cholesterol concentration, but that *T*_m_ is too low to be accessible in our simulations.

When the concentration is increased from 10% to 30%, the lateral diffusion constant reduces by a factor 2 to 3 (see Fig. S3), while viscosity increases by a similar amount (factor of 3 to 4, see Fig 2 (C)). While the values of viscosity observed here (∼ 10^−2^ −10^−1^ P) are much lower than previously reported values in (11), they are inline with other simulations of lipid bilayers (32, 33). The underlying reason for this discrepancy is not entirely clear (see SI for further discussion). The change in diffusion with cholesterol was previously reported in biological-like membranes (34) and thus suggests lipid packing as the main determinant of diffusion, virtually independent of composition.

Increasing temperature tends to decrease saturation and nematic order parameter (see Fig. S4). Moreover, it lowers the response of the membrane to changes in cholesterol concentration, for both viscosity and composition. This arises from the mixing entropy, which has a contribution to the free energy proportional to the temperature and tends towards equal concentrations. In biological membranes, fatty acids located on *sn*2 are mostly saturated and the few exceptions involve unsaturated fatty acids on *sn*1 (35). This may be a way to counteract dominance of the mixing free energy as it strongly reduces the available number of poly-unsaturated species.

The molecular composition here does not correspond to a specific animal species, as biology does not go by the equal-binding approximation and tends to suit particular biological needs (36). However, we can still infer *general* trends in the mechanism, with the caveat that membrane composition of a particular species will be more complex than our model. In fact, our results closely resemble experimental observations in zebrafish (19), where lipids with no or only one saturated tails become more prevalent at higher temperatures, along with cholesterol concentration. It is also reminiscent of trends observed along the secretory pathway, where cholesterol concentration increases (9) along with acyl tail saturation (10).

It is likely that the demixing temperature *T*_m_ is influenced by embedded membrane proteins, which are not modeled here. Some of these are known to bind preferentially to specific lipids (24, 37) and induce domain formation, while others bind to the actin network, which is hypothesized to be involved in membrane structure regulation (38–40). Nevertheless, the increase in nematic order correlation length suggests that *T*_m_ increases with cholesterol fraction, in opposition to model membranes.

The homeoviscous hypothesis states that biological membrane viscosity is regulated and kept constant by cells. Since viscosity is normally inversely proportional to temperature, our results suggest that cells need to increase their cholesterol concentration to compensate for temperature effects. This provides a rationale for the trends observed in zebrafish (19). In fact, viscosity changes rapidly at plasma membrane physiological cholesterol concentrations and our model predicts that at such concentrations, a change of ≈ 5 − 10% cholesterol is required to keep viscosity constant when temperature is changed by ≈ 12 K. This is similar to the change of 4% cholesterol fraction for an 8 K temperature change observed in (19).

## CONCLUSIONS

The regulated ensembles presented in this paper provide a new picture of acyl tail regulation: lipid saturation is a consequence of the other components, such as cholesterol, and is not strictly regulated. It provides a justification for cholesterol trends observed in zebrafish (19), in which increases in cholesterol cause a decrease of acyl tail saturation as well as increased viscosity and correlation lengths of the membrane. In this picture, cholesterol is essentially responsible for regulation of both viscosity and *T*_m_.

The present framework offers many opportunities to further improve the description of complex biological cellular processes. The equal-binding approximation, while a more faithful description than usual ternary model membranes, could be improved to better account for the relevant biology. More work is needed in order to better approximate the chemical networks involved, in particular to understand which ones can be easily extracted into SGC and how to better approximate chemical potential differences between lipid species in order to lift the equal-binding approximation. While there is considerable overlap between lipids chosen here and those found in biological membranes, there are discrepancies between the two sets. Notably, our lipids allow too many unsaturations on *sn*2 and only a single acyl chain length on *sn*1. Better lipid sets, closer available lipids in cells, will be tackled in future research, along with the modeling of asymmetric membranes. In the mean time, we hope that further experiments can lead to more insights into a chemical network based regulation mechanism and provide concrete evidence into the RTE.

## AUTHOR CONTRIBUTIONS

M.G. and T.B. designed the research. M.G. carried out all simulations, analyzed the data. M.G. and T.B. wrote the article.

## ACKNOWLEDGMENTS

We acknowledge Bernadette Mohr, Joseph Rudzinski and Alessia Centi for critical review of this manuscript and Burkhard Dünweg for insightful discussions. This project was supported by the Deutsche Forschungsgemeinschaft (DFG) and the Alexander von Humboldt-stiftung (AvH). This work used computational resources from the Max Planck Computing and Data Facility (MPCDF).

## SOFTWARE

Simulations were run in the HOOMD-blue molecular dynamics engine (41, 42). Systems were assembled using the hoobas molecular builder (43). Simulations are run using a slight modification to the MARTINI force-field (44)(see SI). Visualization was done using the Ovito package (45). Data analysis was done using custom C++ code on top of publicly available C gsd API.

